# The fitness costs of reproductive specialization scale inversely with organismal size

**DOI:** 10.64898/2025.12.08.693006

**Authors:** Christopher Zhang, Eric Libby, Anthony Burnetti, Matthew Herron, William Ratcliff

## Abstract

The evolution of reproductive specialization represents a fundamental innovation in multicellular life, yet the conditions favoring its evolution remain poorly understood. Here, we develop a population genetic framework that examines the fitness cost of reproductive specialization as a function of organism size. We show analytically that the costs of specialization decrease dramatically with organism size. For example, while a 4-cell organism with 50% somatic cells experiences a 50% reduction in population-level exponential growth rate (the same as the two-fold cost of sex), a hundred-cell organism faces only a 15% reduction, a thousand-cell organism 10%, a million-cell organism 5%, and a billion-cell organism merely 3.3%. This scaling relationship arises from the fact that proportionally more cellular growth in larger organisms is required for development, reducing the rate at which the fitness costs of specialization are compounded over multicellular generations. We contextualize our mathematical model with data from the volvocine green algae, showing that simple theoretical predictions closely match empirical measurements. While cellular differentiation demands that somatic advantages compensate for lost reproductive potential, we demonstrate that these compensatory requirements diminish with the logarithm of organism size, fundamentally altering the cost-benefit landscape for large organisms and potentially driving the evolution of a size-differentiation ratchet. This size-scaling relationship helps explain the broad association between large organismal size and multicellular complexity.

## Main

Germ-soma differentiation stands as one of the most consequential innovations in the evolution of complex life (1, 2). The separation of reproductive and somatic functions enabled the morphological diversity characterizing ‘complex’ multicellular life (e.g., animals, plants, fungi, red algae and brown algae). Yet this innovation imposes a fundamental cost: somatic cells, by definition, forfeit reproduction to perform other functions.

Many evolutionary innovations impose costs on exponential growth rates while providing compensating benefits. The two-fold cost of males is the canonical example (3). Sexual populations grow at half the rate of asexual populations because males are incapable of directly generating offspring. This cost persists across diverse taxa, requiring substantial benefits to maintain sex (4). Bacteria face a similar trade-off over metabolic investments, where growth rate depends critically on the proportion of the proteome that goes towards ribosome production (5). The upper bound on cellular growth rate is complete investment in ribosomes, as ribosomes are required to make more ribosomes, generating exponential growth in biosynthetic capacity. Of course, a cell must contain proteins other than ribosomes, but such investment comes at a cost to maximum growth rate. These examples demonstrate a key principle: maximum growth rates are achieved when replicators produce more replicators, and investment in non-reproductive functions is only evolutionarily stable when it provides sufficient compensating benefits.

We apply this framework to understand the evolution of reproductive specialization, often referred to as germ-soma differentiation. In multicellular organisms with specialized cell types, somatic cells enhance survival, resource acquisition, or competitive ability, but are not eligible to create propagules. At the end of an organism’s lifetime, these cells are discarded, and represent wasted potential reproduction (6). The severity of this reproductive cost should constrain which organisms can afford extensive cellular differentiation.

Empirically, the extent of cellular differentiation varies systematically with organism size. Larger species typically have more cell types (7), and the evolution of germ-soma differentiation in volvocine algae follows this pattern clearly: cellular differentiation is absent in small species like *Gonium* and *Pandorina*, while larger species like the 2,048 celled *Volvox carteri* contain less than 1% germ cells (8). The evolutionary basis for this association remains unclear. One possibility is that the fitness costs of somatic specialization decrease with organism size, making extreme specialization prohibitively expensive for small organisms while permitting it in large ones. To investigate this possibility, we develop a simple model of multicellular life cycles, focusing on organisms like the volvocine green algae that develop clonally from single cells, then release germ cells to initiate the next generation (8). In such organisms, only germ cells contribute to the next generation, while somatic cells are discarded at reproduction. This creates an inherent cost: organisms with more somatic cells produce fewer offspring, reducing their population growth rate in the same way that males reduce the growth rate of sexual populations (Figure 1). Our model, developed below, reveals that while investment in soma imposes substantial costs on small organisms, these costs decrease logarithmically with organism size, with larger organisms experiencing proportionally smaller reductions in growth rate for the same investment in somatic cells.

**Fig. 1.**
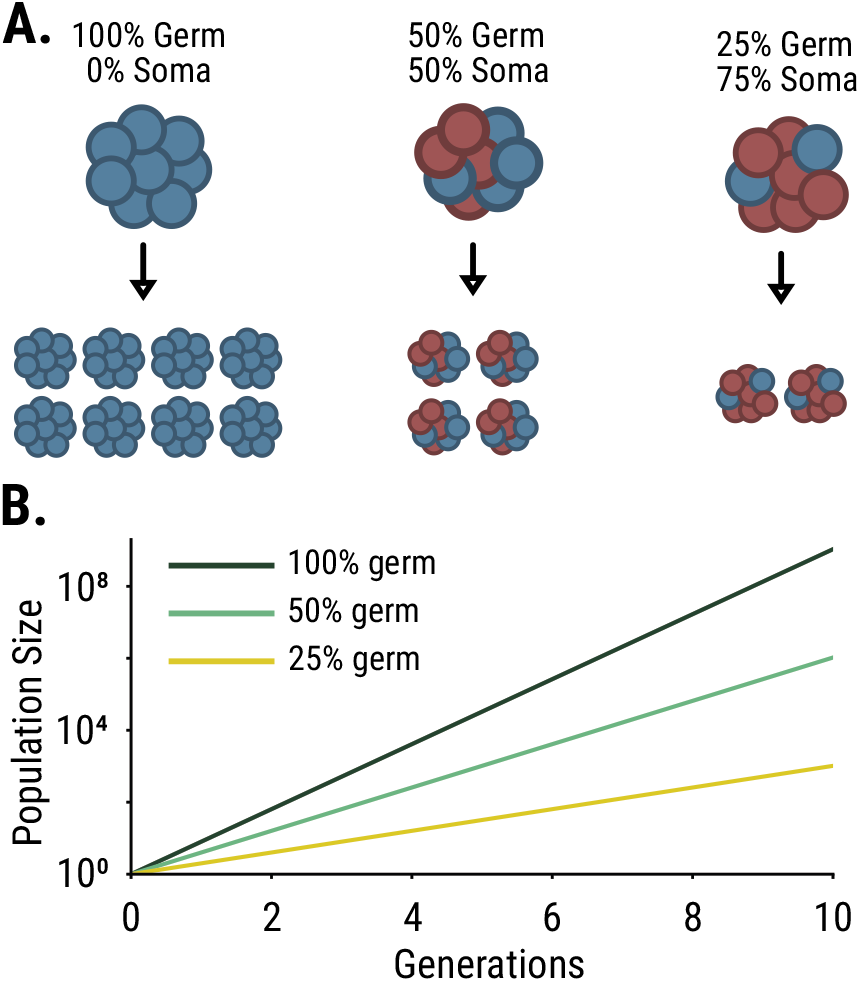
Germ-soma differentiation reduces maximum population growth rates. A) Schematic showing multicellular organisms with different proportions of germ cells (blue) and somatic cells (red). At reproduction, only germ cells contribute to the next generation, while somatic cells are discarded. B) Population growth trajectories over 10 generations for organisms with 100% germ cells (blue), 50% germ cells (green), and 25% germ cells (red). Like the ‘two-fold cost of males’, a 50% allocation to germ cells reduces growth rates by half, absent compensating benefits.

Consider organisms developing from unicellular propagules through synchronized cell divisions to size *N*, with proportion *p*_*g*_ as germ cells. Development requires *τ* = log_2_(*N*) cellular generations. At maturity, the somatic cells die, *N*·*p*_*g*_ germ cells disperse as propagules, initiating the next generation (Figure 2A). Note that this model is agnostic to the timing of germ-soma differentiation: the fitness costs depend only on the final proportion of germ cells at maturity, not on whether differentiation occurs early or late in development, so long as somatic cells continue to divide during development. Population size after *n* generations (time *nτ*) equals (*N*·*p*_*g*_)^*n*^. For continuous time *P* (*t*) = *P* (0)(*N*·*p*_*g*_)^*t/τ*^, where *P* (0) is the initial population size.

**Fig. 2.**
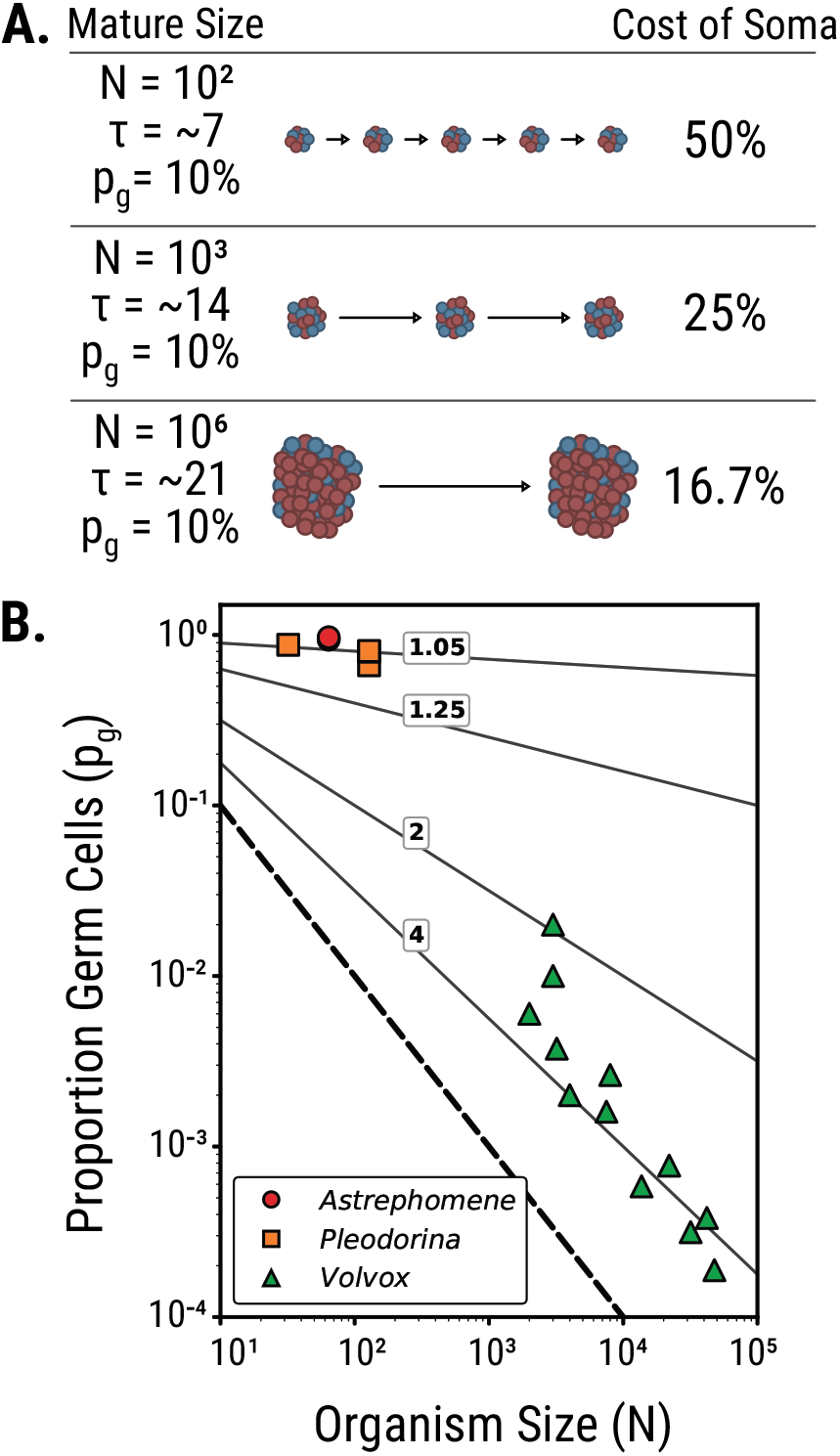
Large organism size reduces the fitness costs of somatic specialization. **A)** Schematic showing multicellular development from a unicellular propagule through synchronized cell divisions to mature size N, with blue cells representing germ cells and red cells representing somatic cells, *p*_*g*_ = the proportion of germ cells. Development requires *τ* = log_2_(*N*) cellular generations. At maturity, only germ cells contribute to the next generation while somatic cells are discarded. The cost of somatic specialization (measured as reduction in growth rate) decreases dramatically with organism size. For organisms with 10% germ cells, a 100-cell organism experiences 50% growth reduction, a 1,000-cell organism faces 25% reduction, and a million-cell organism experiences only 16.7% reduction. Over the same number of cellular generations, larger organisms undergo fewer reproductive events leading to a reduced proportional somatic cost. **B)** Contours show the fold reduction in the per-generation exponential growth rate as a function of organism size for different proportions of germ cells. Larger organisms experience diminished growth rate penalties from somatic specialization. The dotted line represents the theoretical minimum where organisms maintain exactly one germ cell (costs asymptote to infinity at this line). Colored symbols show the calculated growth-rate trade-offs experienced by different species of volvocine algae; the data contained within this plot is enumerated in the supplementary information file and provided code.

The intrinsic growth rate *r* satisfies *P* (*t*) = *P* (0) *e*^*rt*^. Solving yields:

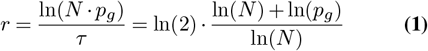

Maximum growth rate occurs when all cells are germ cells (*p*_*g*_ = 1), giving *r*_max_ = ln(2) independent of organism size. The cost of somatic specialization equals:

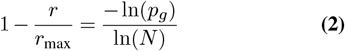

This formula demonstrates a key insight: somatic specialization becomes progressively less expensive as organisms grow larger. Suppose an organism allocates 90% of its cells towards soma, so that *p*_*g*_ = 0.1. The cost of this somatic specialization is 50% in a 100-cell organism, but drops to 25% in a 10,000-cell organism and only 16.7% in a million-cell organism (Figure 2B).

This size-tempering effect arises from the relationship between development time and the geometric increase caused by multicellular reproduction. Small organisms complete development in few cell divisions, causing reproductive losses to compound frequently. Larger organisms require more cell divisions per reproductive cycle, with more biomass doublings between generation transitions. Since development time scales as log(N) while reproductive losses are a fixed proportion of total cell number, larger organisms dilute the rate at which the costs of reproductive specialization are compounded across longer developmental windows, reducing its impact on fitness.

The volvocine green algae provide a natural test of this scaling relationship. In this group, larger organisms have evolved progressively greater somatic investment (Figure 2C; (9)). We quantified this pattern using linear regression on log-transformed species-level data (maximum reported cell number and gonidia count per species). Across 18 volvocine species with germ-soma differentiation, the proportion of germ cells scales strongly with organism size, following *p*_*g*_ ∝*N*^−1.29^ (*r*^2^ = 0.97, *p <* 10^−12^). This relationship holds within the genus *Volvox* alone (*p*_*g*_ ∝*N*^−1.19^, *r*^2^ = 0.85, *p <* 0.0001, *n* = 12 species), spanning organisms from 2,000 to nearly 50,000 cells. Our model predicts that *Volvox aureus*, with approximately 2,040 somatic cells and eight germ cells (typical for this species; 10, 11), should grow 3.7-fold slower than it would if every cell were a germ cell. This is similar to the observed 3-4 fold difference in growth rates between *Volvox aureus* and its unicellular relative *Chlamydomonas reinhardtii* (12, 13).

The analysis above demonstrates that the fitness costs of reproductive specialization scale inversely with organism size. To understand how costs and benefits jointly shape evolution, we must consider what benefits somatic cells provide. While the specific benefits vary across taxa—somatic cells may enhance locomotion, provide structural support, facilitate nutrient transport, or improve survival in other ways—we can explore the general principle through an illustrative model.

We consider the case in which somatic cells offer a benefit by increasing survival of groups in response to repeated episodes of some stressor. For simplicity we assume that these episodes occur every *t*_*s*_ time units so that within a given time *t* there are *t/t*_*s*_ events. As a consequence of these events, only a fraction of groups survive and this survival depends on the group investment in soma, which is 1 − *p*_*g*_. We define this survival function *s*(*p*_*g*_) so investment in soma provides a simple linear increase in survival:

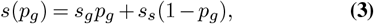

where *s*_*g*_ and *s*_*s*_ represent the survival fraction of a group of only germ or soma cells, respectively.

We can incorporate this survival function into the equation describing the population dynamics of groups to get

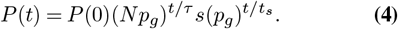

Using this equation, we can determine the investment in germ/soma that maximizes fitness in terms of the population size. We take the derivative with respect to germ investment *p*_*g*_ and find the optimal fitness occurs when 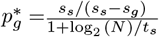. From this equation, we find that there is a critical time 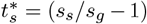 log_2_ (*N*)) for stress events such that when 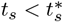 groups that invest in soma are fitter than those that only invest in germ cells (additional calculations in the Supplementary Methods). Interestingly this critical time depends on the size of groups so that larger groups require less frequent events to select for specialization. Thus, for any frequency of stress events there is some group size above which groups increase their fitness by producing somatic cells.

This simple model also reveals a positive feedback dynamic. Once groups invest in soma (*p*_*g*_ *<* 1), increasing in size leads to increased fitness. This occurs even in the absence of stress events because the population equation *P* (*t*) = *P* (0)(*N p*_*g*_)^*t/τ*^ has a positive derivative for 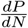 when *p*_*g*_ *<* 1. Indeed, there was nothing about the chosen survival function *s*(*p*_*g*_) that favors larger groups. Rather, the fitness advantage favoring larger groups emerges because smaller groups have shorter life cycles. The cost of investing soma is then not only paid more often over some time period but it compounds geometrically. As groups evolve to be larger the optimal investment in somatic cells also increases, because the optimal 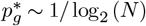 for large *N* .

This feedback dynamic predicts a positive correlation between organism size and investment in cellular specialization: as lineages evolve larger size, they can afford greater somatic investment, which in turn favors further size increases, etc. Consistent with this prediction, larger species have evolved more distinct types of somatic cells (7), and within clades such as the volvocine green algae, size explains the vast majority of variation in somatic investment (Figure 2C). Size is not the only factor shaping cellular differentiation, of course. Previous theoretical work on the volvocine algae showed how clade-specific factors, including nutrient transport and motility constraints, interact with cellular differentiation to shape fitness, and derived the same inverse scaling of specialization costs with organism size that we do (14). Here we demonstrate that this scaling law is a general mathematical consequence of clonal development from unicellular propagules, suggesting it has shaped the evolution of reproductive specialization across the independent origins of ‘complex’ multicellular life.

Most work on the evolution of reproductive specialization has focused on the benefits of division of labor (1, 2, 15, 16). Our results complement this research by quantifying the costs that these benefits must overcome, and showing that these costs depend critically on organism size. By reducing the evolutionary costs of specialization, larger size removes a key constraint limiting reproductive division of labor during evolutionary transitions in individuality.

## Supporting information

Supplemental calculations/data

## Data Availability Statement

All data analyzed during this study are included in this article. Code to reproduce all figures is publicly available at github.com/Sacrozhangt/Size-and-Somatic-Specialization.

## ACKNOWLEDGEMENTS

We thank members of the Ratcliff lab for helpful discussions. This work was supported by John Templeton Foundation award 63580 to WCR and Department of Education GAANN fellowship P200A210046 to CZ. This material is based upon work while MDH was serving at the National Science Foundation.

