## Supplemental calculations/data for "The fitness costs of reproductive specialization scale inversely with organismal size"

### Supplemental Methods

#### Derivation of optimal germ-soma allocation with repeated stress events:

In the main text, we presented a model in which somatic cells provide a survival benefit to multicellular groups facing periodic environmental stress. Here we derive the key results: the optimal proportion of germ cells that maximizes fitness, and the critical frequency of stress events required for somatic specialization to be favored over purely germline investment.

We describe the population dynamics of groups as:

$$P(t) = P(0)(Np_g)^{t/\log_2(N)} \tag{1}$$

Using this we can calculate $\frac{\partial P}{\partial N}$ to determine the group size that maximizes fitness. We find

$$\frac{\partial P}{\partial N} = P(0)(Np_g)^{t/\log_2(N)} \left( \frac{-\log(p_g)\log(2^t)}{N(\log(N))^2} \right) \tag{2}$$

We note that if $p_g = 1$ then the equation has 0 derivative. This mean that fitness cannot be improved by changing group size, since all cells are germ and there is no inherent advantage for groups being big or small. However, if $p_g < 1$ then the derivative is always positive (note $-\log(p_g)$ is positive) so that fitness always increases by increasing $N$. If we consider somatic cells that increase survival then the population dynamic equation becomes

$$P(t) = P(0)(Np_g)^{t/\log_2(N)}(s(p_g))^{t/t_s} \tag{3}$$

We can determine the optimal fitness as a function of $p_g$ by taking the derivative with respect to $p_g$. For simplicity we rewrite the population dynamic equation as:

$$P(t) = P(0)e^{t\left(\log(Np_g)/\log_2(N)+1/t_s\log(s(p_g))\right)} \tag{4}$$

We take the derivative with respect to $p_g$ and find that

$$\frac{\partial P}{\partial p_g} = P(0)e^{t\left(\log(Np_g)/\log_2(N)+1/t_s\log(s(p_g))\right)}\left(1/(\log_2(N)(p_g))+1/t_s(s_g-s_s)/(s_s+p_g(s_g-s_s))\right) \tag{5}$$

We can set this derivative to zero to find the critical value of the $p_g$ that maximizes fitness. Solving for $p_g$ we find $p_g = \frac{s_s/(s_s-s_g)}{1+\log_2(N)/t_s}$. We can find the critical time for stress events $t_s$ by solving this equation for where $p_g < 1$.

DRAFT

#### Volvocine algae data collection:

We compiled species-level data on cell number and gonidia quantity from published literature on volvocine algae. For each species, we recorded the range of reported total cell number and range of number of gonidia per organism from morphological studies. The proportion of germ cells ($p_g$) in Figure 2C was calculated as the ratio of maximum gonidia to maximum total cells. Because volvocine species exhibit variation in cell number, we used maximum reported values to provide a consistent measure of physiological capacity across species. Data on volvocine algae cell numbers and germ cell allocation were compiled from the following sources and are enumerated in supplemental dataset 1.

#### Primary literature used to compile supplemental dataset 1:


1. Janet R. Stein. A Morphological Study of Astrephomene Gubernaculifera and Volvulina Steinii. *American Journal of Botany*, 45(5):388–397, 1958. ISSN 1537-2197. doi: 10.1002/j.1537-2197.1958.tb13142.x. _eprint: https://bsapubs.onlinelibrary.wiley.com/doi/pdf/10.1002/j.1537-2197.1958.tb13142.x.
2. Hisayoshi Nozaki. Morphology and taxonomy of two species of astrephomene in japan. *Journal of Japanese Botany*, 58(11):345–352, 1983. doi: 10.51033/jjapbot.58_11_7555.
3. Hisayoshi Nozaki, Haruko Kuroiwa, Takashi Mita, and Tsuneyoshi Kuroiwa. Pleodorina japonica sp. nov. (Volvocales, Chlorophyta) with bacteria-like endosymbionts. *Phycologia*, June 1989. doi: 10.2216/i0031-8884-28-2-252.1. Publisher: Taylor & Francis.
4. M. O. P. Iyengar and T. V. Desikachary. *Volvocales.* Indian Council of Agricultural Research, New Delhi, 1981.
5. Hisayoshi Nozaki. Notes on microalgae in japan (10). eudorina illinoisensis (chlorophyta, volvocales). *Japanese Journal of Phycology*, 34:143–144, 1986.
6. Gilbert M. Smith. A comparative study of the species of *Volvox*. *Transactions of the American Microscopical Society*, 63(4):265–310, October 1944. doi: 10.2307/3223302.
7. Hisayoshi Nozaki. Morphology, sexual reproduction and taxonomy of Volvox carteri f. kawasakiensis f. nov. (Chlorophyta) from Japan. *Phycologia*, June 1988. doi: 10.2216/i0031-8884-27-2-209.1. Publisher: Taylor & Francis.
8. G. Kochert. Differentiation of reproductive cells in Volvox carteri. *The Journal of Protozoology*, 15(3):438–452, August 1968. ISSN 0022-3921. doi: 10.1111/j.1550-7408.1968.tb02154.x.
9. M.A. Pocock. Volvox and associated algae from kimberley. *Annals of the South African Museum*, 16(3):473–521, August 1933. doi: 10.1111/j.1550-7408.1968.tb02154.x.
